# A maternal effect shapes early-life adaptive body size variation in house mice

**DOI:** 10.64898/2026.08.12.744302

**Authors:** Sylvia M. Durkin, Christina Gao, Michael W. Nachman

## Abstract

Almost all phenotypic variation is mediated by a combination of genetic and environmental effects. Importantly, the balance of these effects can fluctuate over an organism’s lifespan, with the maternal environment being particularly important during early life, especially in mammals. Using house mice as a model, we combine behavioral, molecular, and quantitative genetic approaches to understand how genetic and maternal variation together shape phenotypic divergence in a classic example of morphological adaptation. Temperate house mice are larger than tropical house mice, conforming to Bergmann’s rule. We find that cross-fostering leads to pronounced weight restriction in cold-adapted, large-bodied mice, revealing an important effect of maternal environment. We then describe the molecular underpinnings of genetic and maternal influences on weight by identifying both the genetically- and maternally-controlled differences in gene expression in liver, a key tissue regulating growth. Maternally plastic expression variation is largely controlled in *trans*-, while stable, genetic variation is mediated in *cis-*. Additionally, we link maternally-controlled expression variation in growth-restricted mice to known nutrient deficiency signaling pathways. Finally, we identify candidate loci underlying the genetic basis of adaptive body size divergence from a combination of selection scans in wild populations and *cis*-regulated genes that overlap QTL identified in a mapping panel of temperate and tropical mice. Collectively, these results provide insight into how genetic and environmental forces influence adaptive phenotypic divergence in the context of a critical developmental window.

**SIGNIFICANCE:** Adaptive phenotypes arise from both genetic and environmental influences, yet their unique contributions are rarely resolved within the same natural system. Using locally-adapted house mice that conform to Bergmann’s rule, we experimentally disentangled the genetic and plastic determinants of adaptive body size, focusing on the maternal environment. A maternal effect explains the majority of body weight variation during nursing and is linked to transcriptional changes in nutrient sensing pathways. By separating maternally- and genetically-determined transcriptional variation, we revealed distinct regulatory architectures for plastic and stable expression and identified candidate genes underlying adaptive body size divergence. This work offers broad insight into adaptive evolution and specific details on the mechanistic basis of one of the most widespread ecogeographic patterns in nature.

## INTRODUCTION

Complex traits are mediated by both genetic and environmental factors, and determining the relative contribution of these factors to phenotypic variation is a primary goal across many fields of study, from ecology and evolution to human health and disease. Importantly, the balance of genetic and environmental effects can shift over an organism’s lifespan. For example, the heritability of human height and weight increases from infancy to adulthood, with early childhood being strongly shaped by environmental, as opposed to genetic, factors (Dubois et al. 2012; Jelenkovic et al. 2016). This increased influence of environmental variation in early life is widespread across the tree of life and is often driven by parental effects dominating early development before offspring are independent (Mousseau and Fox 1998; Wolf and Wade 2009). In mammals, the influence of the maternal environment may be particularly important due to the close association between mother and offspring during gestation and lactation. (Mousseau and Fox 1998; Reinhold 2002; Wolf and Wade 2009; Moore et al. 2019). Accurately partitioning environmental and genetic variation enables downstream mechanistic work, such as identifying candidate genes underlying adaptive divergence and understanding the molecular pathways through which environmental variation induces plastic phenotypic change. However, understanding the relative roles of environmental and genetic effects is difficult in systems where experimental manipulation is not feasible.

House mice (*Mus musculus domesticus*) have recently expanded into North and South America within the past 500 years, and in turn have rapidly adapted to diverse climates through changes in morphology, behavior, microbial community composition, and physiology, including potential maternal changes such as nest building and milk content (Barnett and Dickson 1984; Lynch 1992; Phifer-Rixey and Nachman 2015; Phifer-Rixey et al. 2018; Suzuki et al. 2020; Ferris et al. 2021; Agwamba and Nachman 2023; Agwamba et al. 2026). Perhaps the most widely studied phenotypic difference between locally adapted house mice is body size variation. Temperate, cold-adapted house mice are larger than their tropical, warm-adapted counterparts (Phifer-Rixey et al. 2018; Ballinger and Nachman 2022) conforming to Bergmann’s rule. Bergmann’s rule is a well-documented ecogeographic pattern in birds and mammals where animals further from the equator have increased body length and weight, reflecting adaptation to differing climates (Bergmann 1847). Previous work in house mice has found strong evidence for genetic variation mediating body size divergence, particularly though changes in gene regulation (Mack et al. 2018; Phifer-Rixey et al. 2018; Ballinger et al. 2023; Durkin et al. 2024; Gutiérrez-Guerrero et al. 2024). However, these studies all focus on phenotypic variation at a single, adult developmental stage. While body size was not found to be phenotypically plastic post-weaning (Ballinger and Nachman 2022), the contribution of environmental variation during early postnatal development remains unknown, despite this period representing the window during which maternal effects may have their greatest influence on offspring growth (Mousseau and Fox 1998; Wolf and Wade 2009).

Here, we investigate both the genetic and environmental determinants of Bergmann’s rule in locally adapted house mice from temperate and tropical habitats, with a focus on the maternal environment. Specifically, we uncover a maternal effect on body weight during nursing, and we link this to changes in the gut microbiome and altered gene expression in the liver. Next, we partition liver gene expression into genetically- and maternally-controlled changes and use allele-specific expression in reciprocal F1 crosses to describe the regulatory architecture of genetically stable vs. environmentally plastic expression variation. Lastly, we combine quantitative trait locus (QTL) mapping and selection scans in wild populations with our stable, genetically-mediated expression variation to pinpoint candidate loci for adaptive body size divergence. Together, our work identifies a previously unknown source of variation influencing a widely observed phenotypic pattern, giving insight into how genetic and environmental factors together mediate complex, adaptive phenotypes during early development.

## RESULTS AND DISCUSSION

### Cross-fostering influences adaptive body size variation in larger, cold-adapted mice

Under common-garden lab conditions, wild-derived inbred mouse strains from Saratoga Springs, NY, USA (SARA) and Manaus, Brazil (MANA) are known to differ in body weight, with SARA mice being 31.5% larger than MANA at adulthood (Figure 1A-B, SARA mean = 15.5g, MANA mean = 11.8g) (Ballinger and Nachman 2022; Ballinger et al. 2023). Phenotypic variation that persists after many generations in the lab is often assumed to be due to purely genetic variation. However, common-garden conditions cannot control for potential parental influence on phenotype. To understand if variation in the maternal environment influences adaptive body size in this system, we cross-fostered (CF) MANA and SARA mice within 24-hours of birth and measured body weight every 4 days until weaning (21 days, Figure 1C).

**Figure 1:**
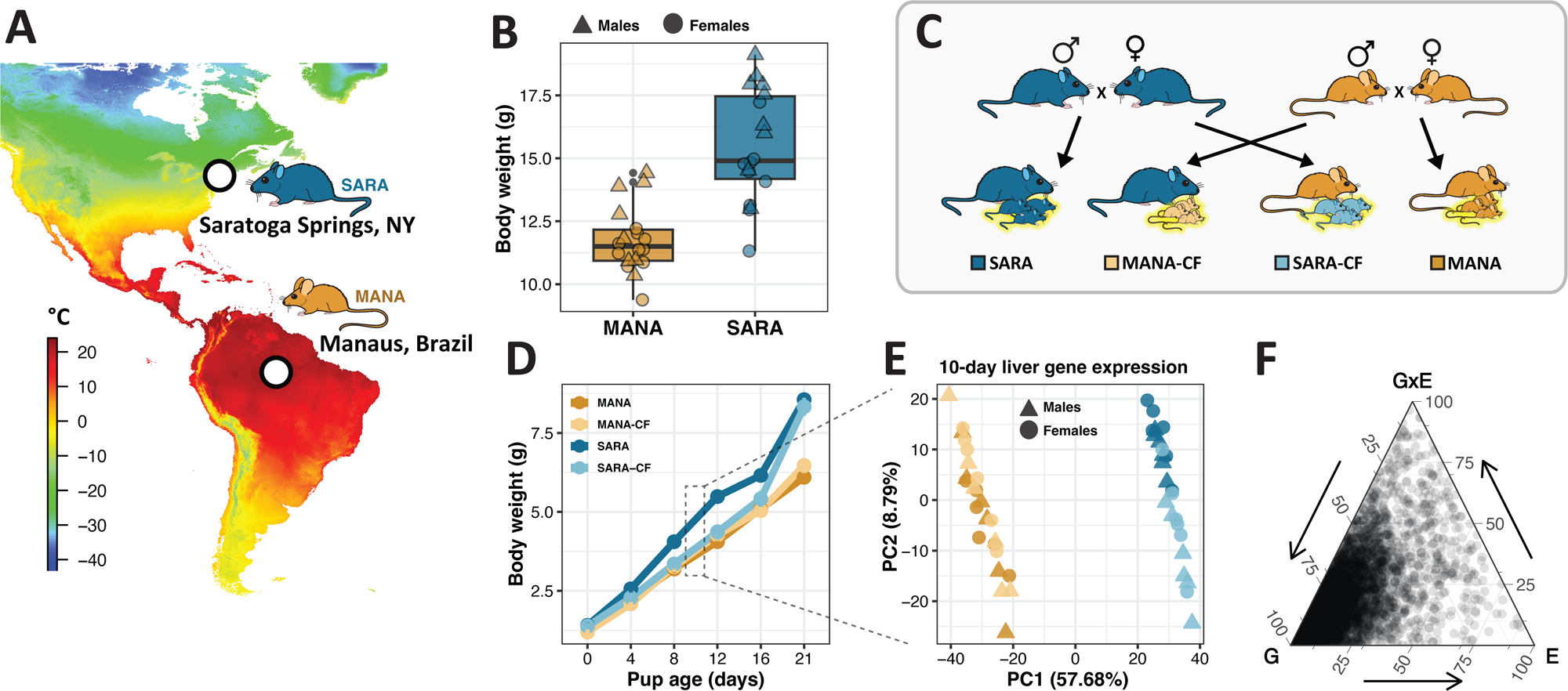
Influence of cross-fostering on body weight and liver gene expression in tropical and temperate house mice. A). Minimum annual temperature at original sampling sites of SARA and MANA lab strains. Panel adapted from Durkin et al. (2024). B). Adult body weight variation in lab-reared SARA and MANA mice. Data are from Ballinger et al. (2023). C). Experimental design of cross-fostering between SARA and MANA mice. D). Body weight variation from birth until weaning (21 days) in cross-fostered (CF) and non-CF pups. (MANA: n=59, MANA-CF: n=22, SARA: n=43, SARA-CF: n=25). E). Principal components plot for liver gene expression from 10-day old CF and non-CF pups. F). Ternary plot showing the proportion of each gene’s expression variation explained by genotype (G), maternal environment (E), and GxE. Each dot represents a single gene and only genes significantly differently expressed in the genotype, maternal, or GxE test are plotted (Details in methods).

There is a strong, unidirectional maternal effect on body weight early in the nursing period where SARA pups reared by a MANA dam are smaller than their non-CF SARA counterparts (Figure 1D; Supplementary Figure 1, p < 0.0001, Welch t-tests). There is no effect of cross-fostering on MANA pups. Interestingly, this weight restriction is mitigated once SARA-CF pups can consume solid food, and not rely solely on the mother’s milk, which happens at about 16 days-of-age (Londei et al. 1988; Silver 1995). At 21 days-of-age, CF pups of both strains are indistinguishable from their non-CF counterparts, and this variation is stable into adulthood (Supplementary Figures 1 and 2). Importantly, we do detect other lasting impacts of cross-fostering, namely gut microbiome composition shifts involving genera with known links to body size variation, which we discuss in later sections.

**Figure 2:**
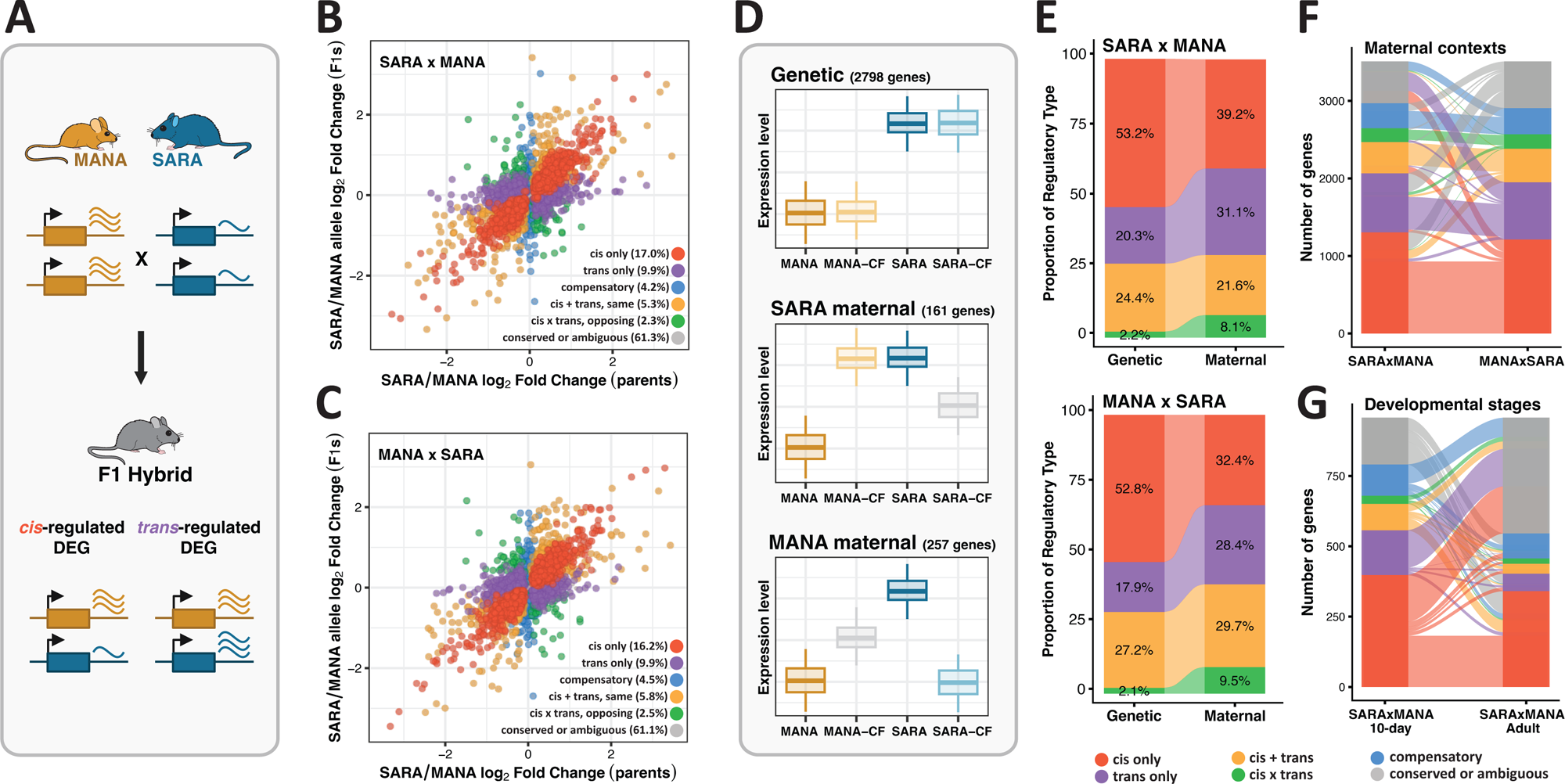
Gene regulatory basis of maternal and genetic expression variation. A). Schematic showing how *cis-* and *trans-*regulatory changes are determined using allele-specific expression (ASE) in F1 hybrids. DEG: differentially expressed gene. B-C). Relationship between parental differential expression and differential expression between alleles in F1 hybrids for SARAxMANA (B) and MANAxSARA (C) F1 hybrid mice. Each dot represents a gene, and genes are colored by regulatory category. D). Schematic of categorizing genetic, SARA maternal, and MANA maternal genes based on relative expression levels, and the number of genes falling into each category. E). Proportion of *cis-* and *trans-*regulation in genetic and maternal categories of genes. Maternal category contains genes in either the MANA maternal or SARA maternal group. Only regulatory types where parental strains show differential expression are included. F-G). Changes in the genes falling within different regulatory categories across maternal contexts (F, SARAxMANA vs. MANAxSARA) and developmental stages (G, 10-day vs. adult SARAxMANA F1 mice).

To understand what may be driving this early postnatal maternal effect on body weight, we analyzed variation in maternal factors including dam weight gain, litter size variation, and care behavior. While SARA dams are larger than MANA dams, within strains there is no difference in weight variation throughout nursing based on whether dams are rearing their birth pups or pups from the opposite strain (Supplementary Figure 3A). Additionally, there is no difference in litter size between strains or between CF and non-CF litters (Supplementary Figure 3B). Lastly, we recorded MANA and SARA dams in their home cage and measured the percent time in the nest at five 1-hour timepoints per day for six days post birth. While MANA and SARA dams partition their time in the nest differently throughout the day, when averaged across all timepoints there is no difference in percent of time spent in the nest (Supplementary Figure 4). While we do not have discrete data on time spent nursing, time in nest can serve as a proxy for the duration of time in which pups could nurse, therefore our data suggest that MANA and SARA pups have equal opportunity to nurse.

Altogether, the lack of divergence in the measured metrics of maternal physiology and behavior suggests that the most likely cause of the maternal effect on weight gain is variation in milk output or composition between SARA and MANA dams that unidirectionally impacts the larger, cold-adapted SARA pups. More explicitly, the smaller MANA dams likely produce less milk or milk of differing content that does not meet the nutritional needs of SARA pups to support normal growth. Importantly, there is evidence that mice adapted to differing thermal regimes vary in milk production and composition (Barnett and Dickson 1984). In wild house mice selectively bred in a cold environment, the milk of cold-adapted dams had higher fat and protein content than control dams, and cold-adapted pups consumed more milk and were larger during the nursing period (Barnett and Dickson 1984). Further, cross-fostering in that experiment showed the same unidirectional effect on weight gain we see here, where cold-adapted pups experienced neonatal weight restriction when reared by control dams, while control pups showed no effect of cross-fostering (Barnett and Dickson 1986).

Neonatal weight restriction due to nutritional deficiency, either in quantity or composition, followed by a sharp increase in weight gain is a well-documented phenomena across mammals, including humans, termed catch-up growth (Hales and Ozanne 2002; Jou et al. 2013; Gat-Yablonski and Phillip 2015). While catch-up growth individuals may not differ from the perspective of overall body weight, there are long-lasting phenotypic impacts that may be costly, such as increased fat deposition, higher incidence of obesity, and greater susceptibility to metabolic disorders, including diabetes (Chen et al. 2011; Jou et al. 2013; Gat-Yablonski and Phillip 2015; Singhal 2017).

### The unidirectional maternal effect on weight gain is recapitulated by gene expression patterns in the liver of 10-day old pups

To understand how the maternal effect on weight gain is reflected at the molecular level, we gathered RNA-seq data at 10-days of age from six male and six female pups from all four groups of CF and non-CF mice, focusing on a metabolically relevant tissue, the liver. We found very little differential expression between sexes at this early developmental stage (Supplementary Tables 1 and 2), and therefore males and females were grouped together in downstream analyses. A principal components analysis (PCA) of liver gene expression variation recapitulates the unidirectional maternal effect on weight gain. Specifically, while PC1 separates samples by strain, PC2 generally separates SARA pups by maternal status, but does not separate MANA and MANA-CF pups, matching the pattern of MANA pups not being affected by cross-fostering at the phenotypic level (Figure 1E).

To further characterize the influence of cross-fostering on expression, we categorized the proportion of individual gene expression variation due to genetic, environmental (maternal), or genotype-by-environment factors (Figure 1F). Consistent with sample clustering along PC1, genotype is the strongest determinant and has a 5.5-times larger contribution to expression variation compared to the maternal environment (measured as the mean absolute value of log_2_ fold change). Overall, while genotype is the main predictor of variation, patterns of liver gene expression in CF and non-CF pups in part match weight gain patterns at the phenotypic level. Because the liver is a key regulator of growth during development, these data present an opportunity to investigate the molecular basis of this maternal effect on body size.

### Maternally-controlled gene expression differences are largely mediated by *trans*-acting factors, while genetic, stable expression variation is largely controlled in *cis-*

In addition to describing the influence of genetic and maternal factors on overall gene expression patterns, we asked *how* this expression variation was mediated at the level of gene regulation using allele-specific expression (ASE) in F1 hybrids. Variation in gene expression can be due to changes in locally acting *cis-*elements (e.g., promoters or enhancers) or longer-range *trans*-acting factors (e.g., transcription factors). *Cis*-changes are thought to be less pleiotropic and more stable across varying conditions as compared to *trans*-changes, which can influence many genes within interconnected networks (Signor and Nuzhdin 2018; Hill et al. 2021; Ballinger et al. 2023; Durkin and Nachman 2025). Due to these factors, *cis*- and *trans*-regulation are predicted to contribute to genetically-controlled and plastic expression variation differently (Smith and Kruglyak 2008; Ballinger et al. 2023). Here, F1 hybrids have one SARA and one MANA allele within a shared *trans*-acting environment. Expression differences between the two alleles can be attributed to *cis*-acting regulation, whereas genes that show no ASE are likely controlled in *trans* (Figure 2A, Wittkopp et al. 2004; McManus et al. 2010; Hill et al. 2021).

In the liver of 10-day old mice, we found that *cis*-regulatory changes predominate differential expression in both SARAxMANA and MANAxSARA F1 hybrids, similar to previous studies in these mice that focused on adult liver expression (Ballinger et al. 2023; Durkin et al. 2024) (Figure 2B-C). Importantly, these overall differential expression patterns between SARA and MANA mice reflect both genetic and maternally-driven expression divergence. To identify both genetically- and maternally-controlled expression and to understand the regulatory architecture of each, we first categorized individual differentially expressed genes (DEGs) based on whether they were insensitive to cross-fostering (Genetic), responsive to cross-fostering by SARA dams (SARA maternal), or responsive to cross-fostering by MANA dams (MANA maternal) (Figure 2D, Supplementary File 1). Unsurprisingly, there are far more genetic DEGs than either maternal set, and more MANA maternal DEGs than SARA maternal DEGs, matching the lack of phenotypic effect of cross-fostering in MANA-CF pups (Figure 2D). The majority of genetic DEGs are controlled in *cis-*, with *cis*-only DEGs being over twice as numerous as *trans*-only DEGs within this gene group (Figure 2E). Conversely, maternal DEGs (either dam) have a significantly higher proportion of *trans*-acting regulation as compared to genetic DEGs (Fisher’s exact test p-value < 0.05 for both SARAxMANA and MANAxSARA comparisons), with the ratio of *cis*-only to *trans-*only genes being more equal (Figure 2E).

Lastly, we investigated the stability of *cis-* and *trans-*regulatory changes across different maternal contexts and developmental stages. To study this across maternal environments, we determined the number of genes that were categorized as a different regulatory type in SARAxMANA hybrids vs. MANAxSARA hybrids (Figure 2F). *Cis*-only genes are more stable across maternal environments (74% shared between comparisons) as compared to *trans*-only genes (56% shared), in agreement with maternally plastic expression having a greater proportion of *trans-*regulation than genetically stable expression. Additionally, we investigated regulatory stability across different developmental stages using previously published liver expression data from adult SARA and MANA mice (Supplementary Figure 5A, Ballinger et al. 2023). While adult liver shows less expression divergence overall as compared to the 10-day data of the present study, *cis*-only genes are far more robust across developmental stages (Figure 2G, 50% of *cis-*changes shared, 8% of *trans*-changes shared), suggesting that the stability of *cis*-regulation applies to both environmental and temporal variation.

In sum, we found that maternally- and genetically-controlled expression differences have distinct regulatory architectures. The importance of *trans*-regulation in mediating plastic responses and the stability of *cis*-changes across environments has been observed in a variety of taxonomic and experimental contexts, including SARA and MANA house mice reared at different temperatures (Ballinger et al. 2023), *Drosophila* species during immune challenge and temperature stress (Chen et al. 2015; Ding et al. 2022), and yeast grown across divergent environmental conditions (Smith and Kruglyak 2008; Tiroshauth et al. 2009). *Trans*-regulatory changes are fast-acting, reversible, and can induce a large suite of coordinated expression changes across a gene regulatory network. These characteristics make *trans*-changes particularly suited to mediating plastic expression variation, despite being more likely to have deleterious pleiotropic impacts (Hill et al. 2021; Vande Zande et al. 2022).

### Maternally-controlled expression variation in SARA-CF mice is dominated by known nutrient deficiency and starvation-induced metabolic pathways

Next, we sought to use the plastic, maternally-controlled DEGs to reveal the molecular basis of the SARA-CF specific weight restriction seen during the neonatal period. To this end, we performed weighted gene co-expression network analysis (WGCNA) on the MANA maternal genes (bottom panel of Figure 2D). Such network-based approaches identify modules of correlated genes that can then be associated with variables of interest, namely maternal status. We identified three total modules; one module was highly correlated with maternal status and contained 66 genes (Pearson’s r = 0.88, p < 0.0001, Supplementary Figure 6). We classified 29 hub genes within this module based on gene expression correlation with 1) module eigengene and 2) the variable of maternal status (Figure 3A).

**Figure 3:**
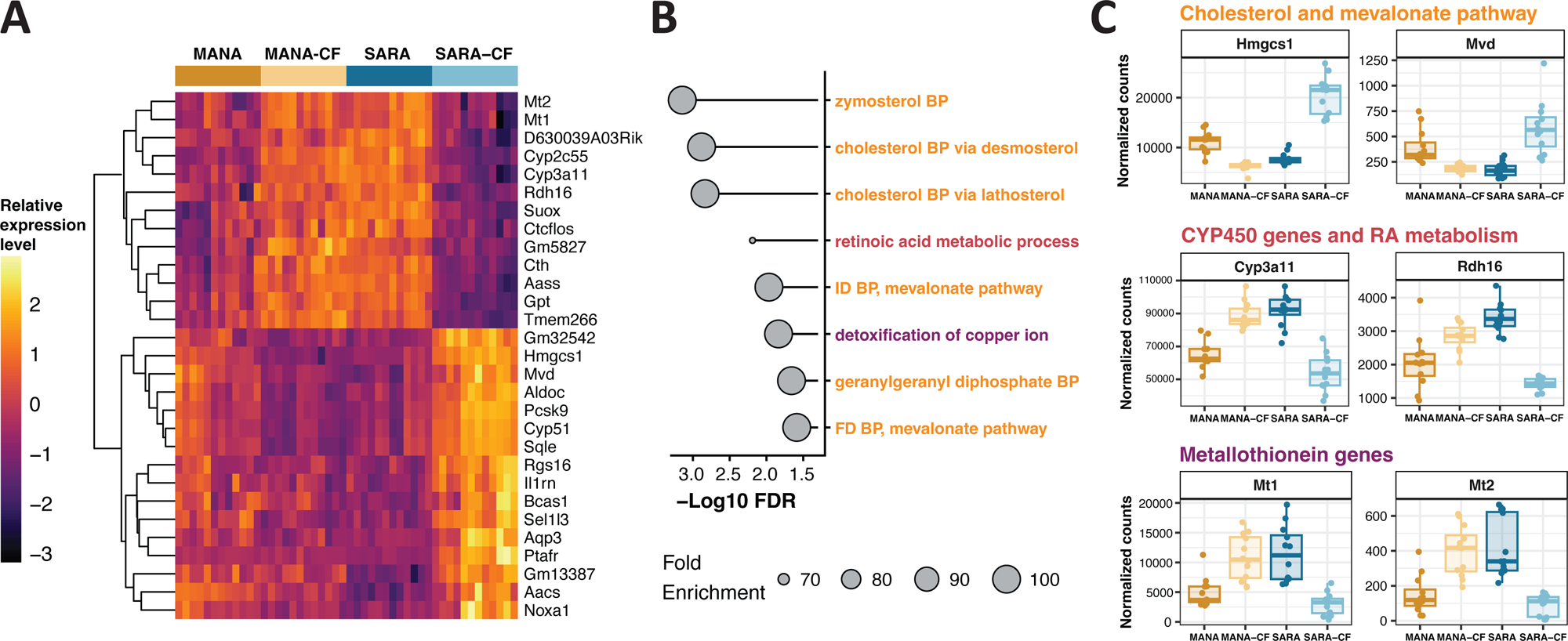
Molecular basis of maternally-controlled weight restriction in SARA-CF mice. A). Hub genes of the WGCNA module most strongly associated with maternal status. Entire module shown in Supplementary Figure 6. B). Gene ontology (GO) enrichment of the 29 hub genes from panel A. Terms are colored based on the specific genes driving the enrichment, as shown in panel C. BP: biosynthetic process, ID: isopentenyl diphosphate, FD: farnesyl diphosphate. C). Expression patterns of a subset of the genes underlying the enriched GO biological processes in panel B.

To understand the biological function of this module, we performed a gene ontology (GO) enrichment analysis on the 29 hub genes. This gene set is enriched for eight biological processes, six of which are related to cholesterol biosynthesis. Other processes point to the importance of retinoic acid (RA) metabolism and detoxification pathways (Figure 3B). The genes underlying the enrichment of cholesterol biosynthesis (*Hmgcs1*, *Mvd*, *Sqle*) are upregulated in SARA-CF mice to a greater degree than all other groups (Figure 3C). *Hmgcs1*, *Mvd*, and *Sqle* encode enzymes in the mevalonate pathway, which is the primary upstream pathway for cholesterol biosynthesis in mammals (Goldstein and Brown 1990). These results point to an activation of cholesterol biosynthesis in SARA-CF mice, potentially in response to nutrient deficiency, explored below.

Cholesterol is a type of lipid, and is an important fuel source in mammalian milk (Görs et al. 2009). Lipid and overall caloric restriction in neonatal mice is known to lead to severe growth restriction (Preidis et al. 2013; Li et al. 2023). When cells are low in lipids, such as cholesterol and fatty-acids, this triggers the upregulation of a family of transcription factors called sterol regulatory element-binding proteins (SREBPs), leading to increased transcription of cholesterol biosynthesis pathway genes (Shimano 2001; Horton et al. 2002). In mice, SREBPs are encoded by the genes *Srebf1* and *Srebf2*, with *Srebf2* being involved in cholesterol biosynthesis (Ye and DeBose-Boyd 2011). Both genes are differentially expressed between MANA and SARA, but only *Srebf2* is under maternal control (Supplementary Figure 7). Further, *Srebf2* was found to be differentially expressed in the liver of other wild-derived house mouse strains in response to diet modulation (Mack et al. 2025).

The genes underlying the enrichments of RA metabolism (*Cyp3a11, Rdh16, Cyp2c55*) and detoxification (*Mt1, Mt2*) are downregulated in SARA-CF mice (Figure 3C). Studies of fasted mice show a decrease in RA biosynthesis and signaling, including retinol dehydrogenase genes (*Rdh* genes) and cytochrome P450 genes (CYP450 genes) (Klyuyeva et al. 2021). Additionally, malnutrition in rodent models is known to lead to decreased CYP450 activity, including CYP3A and CYP2C families (Cho et al. 1999; Zhang et al. 1999; Joo et al. 2004). Lastly, both *Cyp3a11* and *Cyp2c55* are differentially expressed in the wild-derived house mouse diet study mentioned above (Mack et al. 2025), and *Cyp3a11* specifically is known to be involved in weight fluctuations as a function of diet change in mice (Sun et al. 2024).

The last group of enriched genes, *Mt1* and *Mt2*, are metallothionein genes which have been linked to nutritional status in rodents (Szrok et al. 2016) and metabolic disorders, such as obesity and diabetes (Jimenez Jimenez et al. 2025). The primary role of *Mt1* and *Mt2* is to regulate intracellular zinc levels, with a secondary role of regulating copper ions (Chen et al. 2024). Zinc is essential for neonatal growth, with zinc deficient offspring showing growth restriction in both mouse models and humans (Beach et al. 1980; Brion et al. 2020). Importantly, mouse pups receiving milk low in zinc exhibit decreased liver *Mt1* and *Mt2* expression (Vruwink et al. 1988). Overall, maternally-controlled expression variation in SARA-CF pups points to altered regulation of signaling pathways involved in key metabolic processes, and this dysregulation may mediate the weight restriction seen in this experimental group.

### Rearing mother has a lasting influence on the pup microbiome, even after the maternal effect on weight gain is no longer detectable

Because catch-up growth can have lasting physiological impacts in the absence of lasting differences in body weight, we sought to categorize the influence of cross-fostering on an important element of neonatal metabolism, the microbiome. Across mammals, gut microbiota are known to influence a variety of host phenotypes related to ultimate fitness, including body size (Turnbaugh et al. 2006; McFall-Ngai et al. 2013; Yun et al. 2017), and microbiome composition is strongly influenced by rearing mother, often to a greater extent than birth mother (Pantoja-Feliciano et al. 2013; Daft et al. 2015; Treichel et al. 2019). Further, previous work on MANA and SARA, among additional wild-derived mouse strains, suggests that gut microbiota variation may play a role in body weight variation both in the wild and the lab (Moeller et al. 2018; Suzuki et al. 2020).

To investigate gut microbial variation in CF and non-CF pups we performed long-read 16s rRNA sequencing from fecal samples collected at three weeks-of-age, when the microbiome is considered somewhat stable in mice (Pantoja-Feliciano et al. 2013). We also collected samples from the rearing dams of each group, and classified microbial diversity across all samples at the genus level (Figure 4A, Supplementary File 2). In a PERMANOVA analysis of Bray-Curtis dissimilarities, we found that both rearing dam and strain are significant predictors of pup microbial composition, but rearing dam explains 45% (p < 0.01) of variance, while strain explains only 13% (p < 0.05). Additionally, pairwise comparisons showed that the microbial communities of all pup groups are significantly different from one another, except MANA vs. SARA-CF pups (MANA-reared pups) (Figure 4B). Importantly, there was no variation between pup samples for alpha diversity metrics, including Shannon’s and Simpson’s indices (pairwise ANOVA, p > 0.05 for all comparisons), suggesting that pup microbiomes do not differ in the number or evenness of genera present, but instead differ in composition.

**Figure 4:**
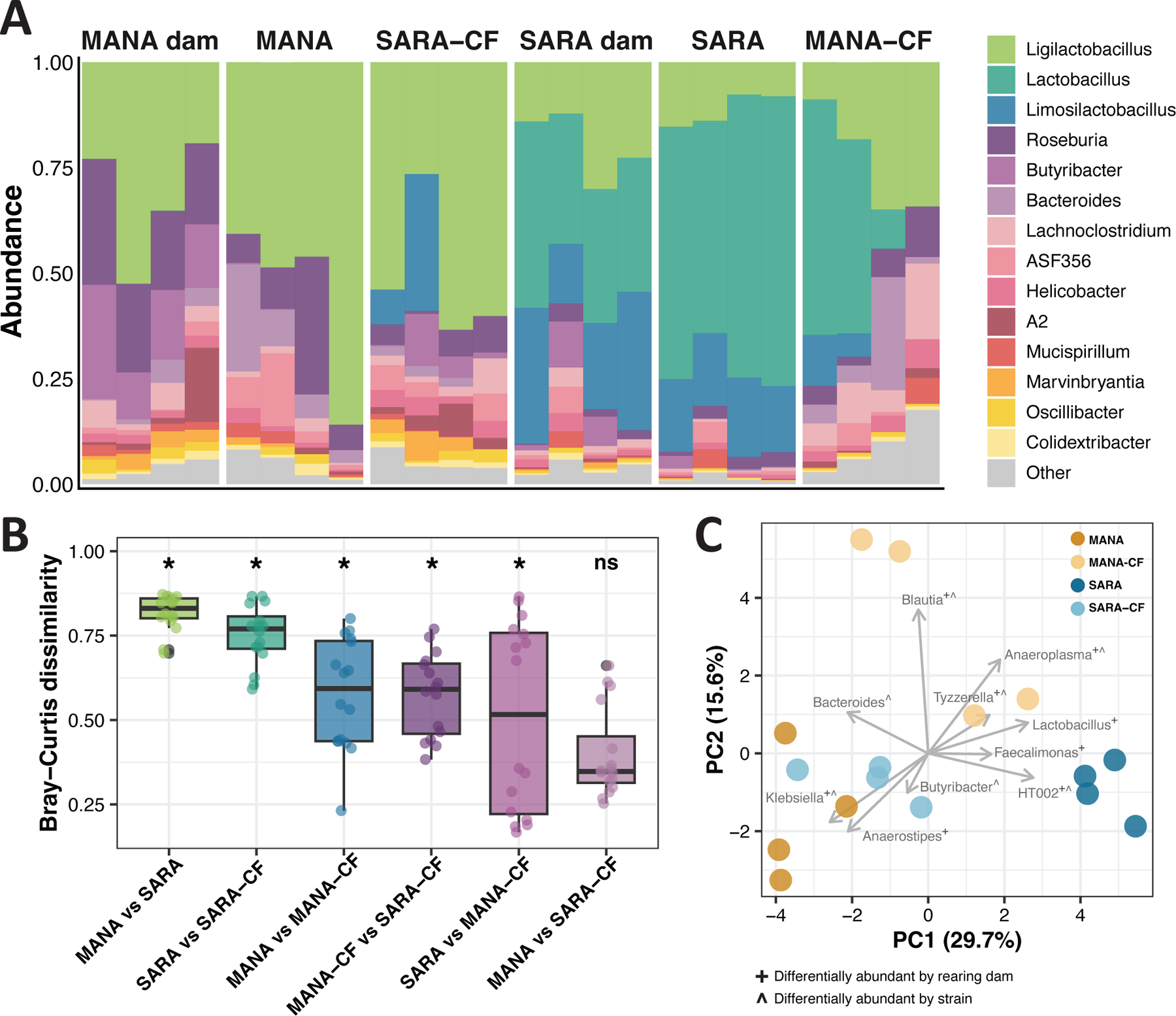
The gut microbiome is strongly shaped by rearing dam. A). Microbiome composition of the 15 most abundant bacterial genera across all dam and pup samples (3-weeks of age). Relative abundances of all detected genera are listed in Supplementary File 2. B). Pairwise Bray-Curtis dissimilarity distributions for each pup group comparison. Comparisons are ordered from most divergent to least divergent. Significant dissimilarity was calculated though pairwise PERMANOVA tests with a Benjamini-Hochberg correction. * p < 0.05. C). Principal components plot of relative genera abundance for all pup microbiome samples. Differential abundances for rearing dam and strain calculated using DESeq2.

To understand the specific genera driving the observed differences in microbial composition, we defined differentially abundant genera both by strain (7 genera differentially abundant) and rearing dam (8 genera differentially abundant) and asked how they related to pup sample clustering in a principal components analysis (Figure 4C). Perhaps the most striking difference in microbial composition is that of *Lactobacillus*, which is highly abundant in SARA dams and SARA-reared pups, while absent in MANA dams and the pups they rear (Figure 4A).

*Lactobacillus* is a prominent component of the neonatal gut microbiome across mammals, and is known to be transferred through milk (Pantoja-Feliciano et al. 2013; Murphy et al. 2017). Importantly, the relative abundance of *Lactobacillus* has links to neonatal weight gain in mice, including promoting weight gain (Schwarzer et al. 2016) and responding to accelerated postnatal growth (Wang et al. 2016). Lastly, there is a connection between *Lactobacillus* abundance and cross-fostering specifically, with the microbiome of pups reared by a high-fat-diet dam having increased *Lactobacillus* relative to littermates reared by a normal-diet dam (Xue et al. 2022). This suggests that the lack of *Lactobacillus* in the microbiome of SARA-CF pups could contribute to the observed weight restriction. Ultimately, the gut microbiome in SARA and MANA pups is strongly shaped by rearing dam, and there are significant shifts in particular genera that may be linked to elements of pup growth.

### QTL regions for body size, *cis*-DEGs, and scans for selection in wild populations reveal candidate genes underlying body weight divergence between SARA and MANA mice

Finally, we sought to describe the genetic basis of body weight by utilizing a published QTL mapping experiment between MANA and SARA mice that identified five genomic regions associated with body weight variation (Durkin et al. 2026). We first narrowed down candidate loci using genes with a known impact on body size, as defined by the Mammalian Phenotype (MP) browser in the Mouse Genome Informatics database (abnormal body size: MP code 0003956, Baldarelli et al. 2024). We found that genes above the logarithm of odds (LOD) significance threshold in the QTL study are enriched for the MP body size genes, as compared to randomly placed markers (Figure 5A, p < 0.01), suggesting that using overlapping QTL and body size MP genes may be fruitful in identifying potentially causal genetic variants underlying weight variation in SARA and MANA mice.

**Figure 5:**
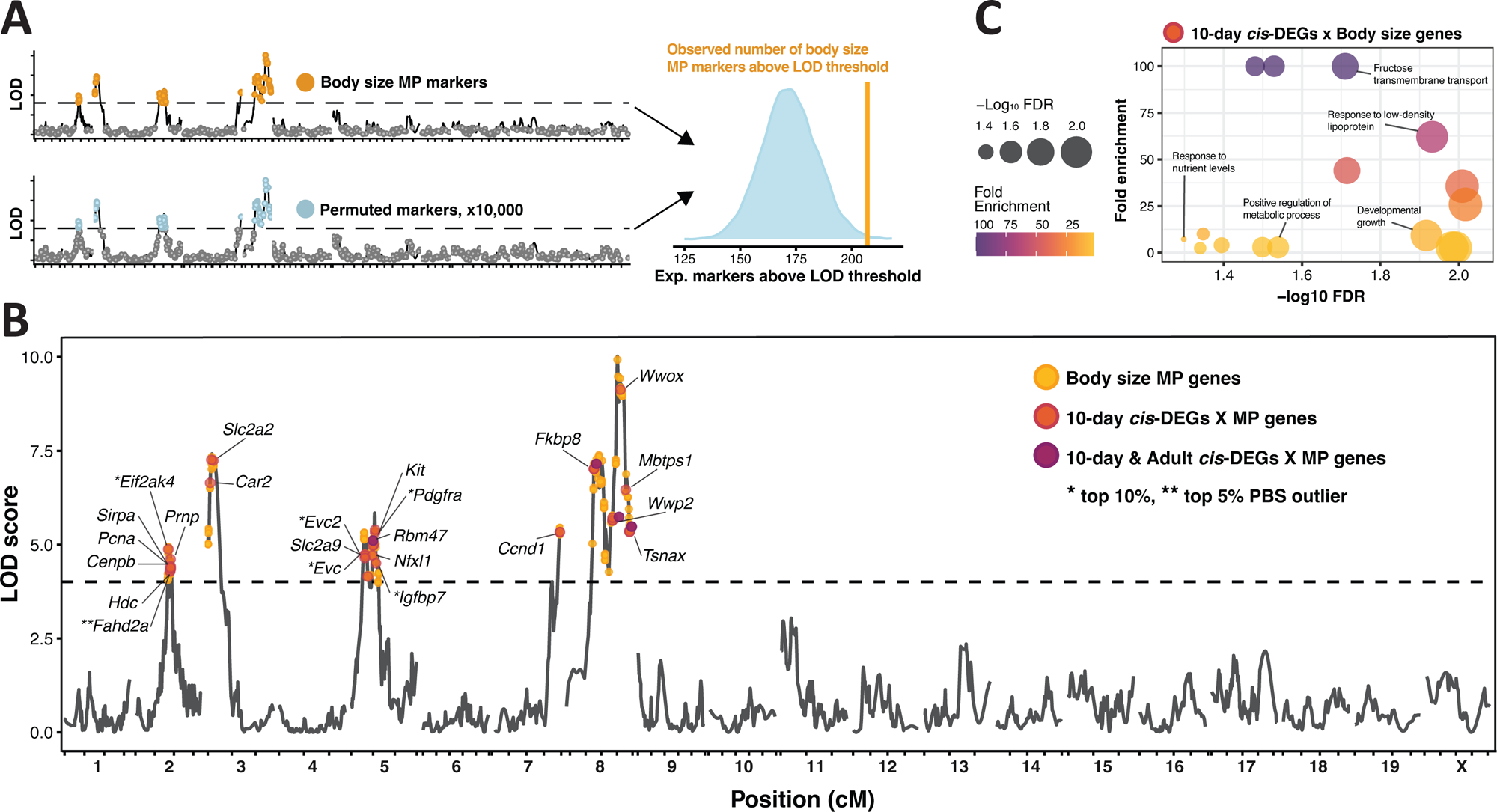
*cis-*regulation associated with adaptive body size variation. A). Schematic of permutation analysis and observed enrichment of body size MP genes above the LOD threshold for body weight QTL. The orange line represents the observed number of body size MP genes above the LOD threshold, and the blue distribution represents the permuted number of genes above the LOD threshold among 10,000 random sets of the same size. B). Plot of genome-wide LOD scores for body weight from Durkin et al. 2026. The points represent the closest markers to genes of interest. Genes that were found to be *cis*-regulated in both the SARAxMANA and MANAxSARA F1 hybrids are labeled. All overlapping *cis*-DEG x MP code genes are listed in Supplementary File 3. C). Gene ontology (GO) enrichment of the 32 *cis*-DEGs (*cis*-regulated in either F1 hybrid) that overlapped body size MP genes within QTL intervals. Full list of terms in Supplementary File 4.

Next, we focused on *cis-*regulated DEGs, as *cis*-regulatory elements are physically close to the genes they regulate and are highly stable across environmental contexts (Figure 2F-G). We looked for overlap between *cis*-DEGs in the liver of 10-day old mice and MP body size genes within QTL intervals (32 genes), as well as *cis*-DEGs found in the adult liver data (4 genes, Figure 5B, Supplementary File 3). In a GO analysis, the 32 overlap candidate genes are enriched for biological processes related to developmental growth, sugar transport, nutrition, and metabolism regulation (Figure 5C, Supplementary File 4). For example, the highest enriched term, fructose transmembrane transport, is driven by the genes *Slc2a2* and *Slc2a9*. These genes are involved in energy homeostasis, with mutations and knock-out phenotypes impacting metabolic diseases such as diabetes and fatty liver disease (Debosch et al. 2014; de Souza Cordeiro et al. 2022; Zeng et al. 2022). Importantly, these GO term enrichments are not due to the candidate genes being a subset of body size MP genes, as random subsets of 32 MP body size genes do not return the same degree of significant GO enrichments (Supplementary Figure 8).

Finally, we used published scans for selection in house mice to connect our lab-based candidate gene identification to natural selection acting in wild populations (Durkin et al. 2024). Specifically, we used a normalized version of the population branch statistic (PBSn1), which identifies allele frequency shifts that are unique to one focal population in comparison to two outgroup populations. We used two published PBSn1 tests, one where Manaus was the focal population, and one where a population on the border of New Hampshire and Vermont (NH/VT) was the focal population. The NH/VT population is geographically close to Saratoga Springs, the original sampling locality of SARA mice. For both tests, France and Iran were outgroup populations, which are near the ancestral range of *Mus musculus domesticus* (Phifer-Rixey and Nachman 2015; Ferris et al. 2021; Morgan et al. 2022).

Six candidate genes are in the top 5% or 10% of PBS outliers (Figure 5B). These include *Eif2ak,* an amino acid sensor that regulates the starvation response in mammals (Battu et al. 2017), *Igfbp7*, a regulator of insulin and insulin-like growth factors (IGFs) associated with metabolic disease (Evdokimova et al. 2012; Yan et al. 2019), and lastly, *Evc* and *Evc2*, which are positive regulators of skeletal growth through the well-conserved Hedgehog signaling pathway (Ruiz-Perez et al. 2007; Blair et al. 2011). Overall, by integrating scans for selection, *cis*-DEGs, experimental crosses in the lab, and genes with known phenotypic effects, we were able to pinpoint strong candidate genes for adaptive body size variation in tropical and temperate house mice. This type of comprehensive work sets the stage for future functional validation with the goal of fully linking genotype to phenotype for naturally occurring complex traits.

## CONCLUSION

Describing how genetic and environmental forces together mediate phenotypic variation is necessary for understanding the process of adaptation. Here, we decompose the contribution of genetic variation and the maternal environment to a widespread ecogeographic pattern observed across animals, Bergmann’s rule, and show how these contributions vary over early postnatal development. We found that cross-fostering strongly influences body weight in cold-adapted, large-bodied mice early in life, but this effect is mitigated once pups can eat food on their own, pointing to a potential role of milk variation in mediating this maternal effect. Strikingly, this unidirectional maternal effect is recapitulated in the gene expression patterns of the liver, a metabolically relevant tissue. We then describe the molecular and gene-regulatory basis of both genetic and maternally plastic expression variation, demonstrating the contrasting roles of *cis-* and *trans-*acting regulation in mediating genetic and maternal expression, respectively, and the induction of canonical nutrient deficiency pathways among maternally-controlled genes. Further, while cross-fostering may not impact body weight past weaning at the phenotypic level, we show altered gut microbial composition as a function of rearing dam, with links to metabolism and weight gain variation later in life. Lastly, the identification of environmentally stable, *cis*-regulated expression allowed us to identify candidate genes underlying the genetic portion of body weight divergence in these mice, showing the utility of investigating both sources of variation in tandem.

Ultimately, these results represent an initial step in fully describing a complex trait such as body size. Further, our findings highlight the importance of an often-overlooked source of variation in lab-based studies, the maternal environment. This integrative work suggests fruitful avenues for future research not only on the genetic basis of body size, but also on the role of the maternal-offspring relationship in shaping environmental adaptation, including variation in milk composition and the long-term impacts of early-life maternal care differences on metabolic and physiological condition.

## MATERIALS AND METHODS

### Ethics statement

This work was conducted in accordance with the University of California, Berkeley Institutional Animal Care and Use Committee (AUP-2016-03-8548-3). Euthanasia was performed under the approval of the ACUC at the University of California, Berkeley, and included the humane use of isoflurane and cervical dislocation by trained personnel.

### Animal husbandry and mice used

All mice were housed under a 12:12 light cycle at 70-72°F and fed standard rodent food *ab libitum* (PicoLab Rodent Diet 20: 20% protein, 4.5% fat, 6% fiber). Wild derived, inbred strains of house mice (*Mus musculus domesticus*) used in this study were originally collected from 43°N in Saratoga Springs, New York (SARA) and from 3°S in Manaus, Brazil (MANA). Information on the establishment of these inbred strains is described in Phifer-Rixey et al. (2018) and Ferris et al. (2021). In brief, live mice were collected from the field, brought back to the lab, and unrelated individuals were mated to create the N1 generation. Inbred strains were generated via subsequent sib-sib mating. SARA and MANA mice used in this study were past generation 20 of sib-sib mating and considered fully inbred.

### Cross-fostering experiment

To assess the presence of postnatal maternal effects on body weight in SARA and MANA mice, we exchanged pups from one strain with pups of the other strain within 24 hours of birth. Whole litters were always exchanged, despite often uneven pup numbers between the switched litters. All litter sizes ranged between 1-6 and do not differ between strains (Supplementary Figure 3B). All experiments were conducted using a dam’s second successful litter to control for litter order effects, which may influence maternal care in mice (Cohen-Salmon 1987; Crusio and Schmitt 1996). We measured body weight of cross-fostered (CF) and non-CF mice at 6 time points throughout the nursing period: PND 0 (postnatal day 0, day of birth), PND 4, PND 8, PND 12, PND 16, PND 21 (weaning). Dam weight was also recorded at each timepoint. Additionally, we measured body weight in a subset of CF and non-CF mice weekly from weaning until 9 weeks of age to track how maternal influence persisted into adulthood. MANA pups that were fostered by a SARA mother are called MANA-CF, and SARA pups that were fostered by a MANA mother are called SARA-CF.

### Maternal behavior observations

To assess variation in maternal care behavior and general activity, we took continuous video recordings of SARA and MANA dams from 6.5 zeitgeber time (ZT, denotes hours since lights on) on PND 1 to 2.5 ZT on PND 7 using GoPro HERO8 4k cameras trained at the home cage with the nest in view. Behaviors were scored at 5 evenly spaced 1-hour intervals throughout the 24-hour day (2-3 ZT, 6.5-7.5 ZT, 11.5-12.5 ZT, 16.5-17.5 ZT, and 21.5-22.5 ZT). Within each hour, behaviors were recorded once every three minutes. In total there were 600 behavioral measurements per mouse: 20 measurements per hour, 5 hours per day, 6 days total. To assess time in nest, we noted whether the mouse was inside or outside the nest at each timepoint.

### RNA extraction, library preparation, and sequencing

At 10 days-of-age, pups were euthanized and liver tissue was collected from six males and six females from the following groups: MANA, MANA-CF, SARA, SARA-CF, SARAxMANA F1 hybrids (SARA dam), and MANAxSARA F1 hybrids (MANA dam). Tissue was immediately placed in RNAlater, kept at 4°C for 24 hours and stored at -80°C until RNA extraction. We extracted RNA using the NEB Monarch Total RNA Miniprep Kit and generated cDNA libraries using the Watchmaker mRNA Library Prep Kit. All libraries were barcoded using unique dual indexes from Illumina, pooled in equimolar ratios, and sequenced across one lane of 150bp paired-end NovaSeq X at the Vincent J. Coates Genomics Sequencing Center at UC Berkeley. Raw reads were trimmed for adapter sequences and filtered for quality (Phred score > 15) using fastp (version 0.19.6, Chen et al., 2018).

### Differential expression analyses

We mapped clean reads to the *Mus musculus* reference genome (GRCm39) using STAR (version 2.7.11, Dobin et al., 2013) and counted mapped reads that overlapped exons in the GRCm39.110 annotation using HTSeq (version 0.11.0, Anders et al., 2015). Separate count files for each sample were merged using the python script merge_tables.py (Dave Wheeler, <u>github</u> <u>link</u>), generating the input for downstream differential expression analyses.

We quantified differential expression between strains and maternal groups using DESeq2 in R (Love et al. 2014). We removed genes with a mean expression level less than 10 reads across samples, resulting in 16,659 expressed genes. We analyzed males and females together as there was very little differential expression between sexes (Supplementary Tables 1 and 2). We computed pairwise differential expression between all strains (MANA and SARA) and maternal groups (CF and non-CF) using Wald tests and the model design: ∼ genotype + maternal + genotype:maternal. We defined a gene as differentially expressed if it had a p-value < 0.05, with Benjamini-Hochberg correction, and a log_2_ fold change value > 0.5 or < -0.5. In addition to our pairwise cohort comparisons, we tested the influence on expression of just genotype (SARA + SARA-CF vs. MANA + MANA-CF) and just maternal type (SARA + MANA-CF vs. MANA + SARA-CF) using Wald tests. Lastly, we quantified GxE effects by contrasting the variation between SARA vs MANA when MANA is the mother as compared to when SARA is the mother. We also used DESeq2 to generate principal component plots.

### Genetic and maternal influence on gene expression

To directly quantify the proportion of liver gene expression variation due to genetic, environmental (maternal), or GxE effects, we first calculated a total variance value for each gene as the sum of the absolute log_2_ fold change values for the genotype test, maternal test, and GxE test (defined above). Proportions were calculated as the absolute log_2_ fold change of each individual effect divided by the total variance.

To further explore the influence of genetic and maternal forces on liver gene expression, we defined sets of genetically- and maternally-controlled genes (Figure 2D). A gene was considered genetically-controlled if it was significantly differentially expression between SARA and MANA and between SARA-CF and MANA-CF, in the same direction. Defining maternally-controlled genes was done separately for MANA and SARA dams. A gene was considered maternally-controlled by SARA dams if it was significantly differentially expressed between SARA and MANA pups, between MANA-CF and MANA pups, and showed the same directionality in differential expression in these comparisons. A gene was considered maternally-controlled by MANA dams if it was significantly differentially expressed between SARA and MANA pups, between SARA-CF and SARA pups, and showed the same directionality in differential expression in these comparisons. All genetic and maternal genes are listed in Supplementary File 1.

### Variant calling for allele-specific expression analyses

We called single nucleotide polymorphisms (SNPs) from MANA, SARA, and F1 hybrids using the genome analysis toolkit (GATK, version 4.2.2.0, Depristo et al. 2011) following the RNA-seq best practices for germline short variant discovery. After aligning, as described above, we prepared BAM files using MarkDuplicates, SplitNCigarReads, and AddOrReplaceReadGroups. We called haplotypes on all individuals separately using HaplotypeCaller, combined resulting gVCFs, and called genotypes jointly using GenotypeGVCFs. We selected SNPs using SelectVariants, filtered for quality using VariantFiltration (QD > 2, FS < 60, MG > 40, MQRankSum > -12.5, ReadPosRandSum > -8), and retained SNPs present in at least 85% of individuals using VCFtools (Danecek et al. 2011).

To assign F1 reads to parental alleles, we first generated a VCF of fixed and different SNPs between MANA and SARA using VCFtools and merged this resulting VCF with SNPs heterozygous in F1 individuals (identified using the GATK VariantFiltration filter “isHet”). This resulted in 63,280 allele-specific expression (ASE) informative SNPs for downstream analyses. Lastly, we phased this VCF to indicate whether SARA or MANA contained the reference allele using a custom script (available on <u>GitHub</u>).

### Allele-specific expression analyses

We separated F1 reads mapping to either MANA or SARA by remapping F1 reads with STAR using WASP (Van De Geijn et al. 2015), specifying the ASE informative SNPs from above as the VCF. WASP adds a SAMtag to each read that indicates its phasing in regard to the provided VCF. We then generated separate MANA and SARA bam files using grep and counted reads using HTSeq as described above. After filtering for genes with a mean count greater than 10 across samples, there were 7,679 and 7,471 genes to analyze for ASE in SARAxMANA and MANAxSARA F1 hybrids, respectively. F1 reads mapped equally to SARA and MANA alleles, suggesting little mapping bias toward either strain (Supplementary Figure 9).

We categorized genes into different regulatory classes using three hierarchical statistical tests of differential expression similarly to McManus et al. (2010). First, we quantified differential expression between MANA and SARA parent samples using Wald tests (P). Second, we tested for differential expression between the alleles of the F1 hybrids using Wald tests (H). Lastly, we tested for the presence of *trans-* effects using a likelihood ratio test comparing parent to hybrid allele expression variation (T). A significant difference between these ratios would suggest that the expression difference between alleles in an F1 does not recapitulate the expression difference between the parents, therefore indicating the presence of *trans*-regulation. All tests were performed using DESeq2 in R (Love et al. 2014). We then categorized genes into different regulatory modes based on their significance at an FDR of 5% in the P, H, and T tests.

- *cis-* only: significant in P and H, not significant in T
- *trans-* only: significant in P, not significant in H, significant in T
- *cis + trans*: significant in P, H and T. Expression divergence direction in P and H is the same.
- *cis* x *trans*: significant in P, H and T. Expression divergence direction in P and H is not the same.
- compensatory: significant in H, not significant in P, significant in T. No expression divergence due to compensation.
- conserved: not significant in H, P, or T. No expression divergence.
- ambiguous: Genes not encapsulated by above parameters.

Genes falling into each regulatory category (excluding ambiguous) are listed in Supplementary File 5.

### Adult liver gene regulatory analyses

To investigate gene regulatory patterns across developmental stages, we integrated the 10-day liver gene regulatory patterns of this study with previously published gene regulatory results from adult MANA and SARA mice (Ballinger et al. 2023). Ballinger et al. (2023) studied gene regulatory patterns in the liver of mice reared at room temperature, and in a cold room (4°C). Starting at the stage of counted reads (available from zenodo doi: 10.5281/zenodo.8288000), we reanalyzed adult liver expression data from MANA, SARA, and SARAxMANA male mice, categorizing genes into different regulatory categories as described above. Mice reared in cold and room temperature conditions were analyzed separately, and the comparison of 10-day expression to the cold room F1 hybrids is shown in Supplementary Figure 5B. Because Ballinger et al. (2023) focused on males, we also redefined regulatory classes from our 10-day SARAxMANA expression data using only males to be comparable. Genes falling into each regulatory category (excluding ambiguous) are listed in Supplementary File 5.

### Weighted gene co-expression network analysis

To isolate groups of genes with expression patterns indicative of maternal control, specifically in weight-restricted SARA-CF pups, we performed weighted gene co-expression network analysis (WGCNA, Langfelder and Horvath 2008) on all MANA maternal genes (Figure 2D, 257 genes). Reads were normalized prior to analysis using the vst function in R/DESeq2 (Love et al. 2014). After determining the appropriate power threshold of 8, we constructed networks using the blockwiseModules function in R/WGCNA specifying TOMType=signed, minModuleSize=20, deepSplit=2, and mergeCutHeight=0.25.

We correlated module eigengene values with variables of rearing dam (MANA vs. SARA) and group (MANA, MANA-CF, SARA, SARA-CF) using the cor function in R/WGCNA to determine which module was most closely related to maternally-controlled expression patterns, with a specific focus on SARA-CF expression. We then defined hub genes within a module similarly to that described by Langfelder and Horvath (2008) by calculating 1) module membership (kME) as the Pearson correlation between each gene’s expression and the module eigengene, and 2) gene significance (GS) as the Pearson correlation between each gene’s expression and the variable of interest (rearing dam or group). Genes with absolute values of kME > 0.8 and GS > 0.4 were considered hub genes.

### Microbiome analyses

We collected fecal samples from 3-week-old pups and their rearing dams using the QIAamp PowerFecal Pro DNA Kit. We collected four samples per pup group (MANA, MANA-CF, SARA, SARA-CF) and two samples per dam group (MANA, MANA-CF, SARA, SARA-CF), for a total of four dams per strain. Each pup group contains samples from at least two litters. Libraries were constructed using the Oxford Nanopore (ONT) 16S Barcoding Kit V14 and sequenced across one ONT PromethION flow cell. Raw pod5 files were converted to fastq format using Dorado’s basecaller function (version 1.4.0) and subsampled to 50,000 reads to match the lowest yield sample using SeqKit (version 2.13.0, Shen et al. 2016). To ensure accurate capture of 16S rRNA reads, we filtered for reads between 1,100 bp and 1700 bp using SeqKit. The average length of reads across all samples was 1,466 bp.

Following fastq processing, we determined the taxonomy of reads and quantified taxa counts with minimap2 (version 2.30, Li 2018) using the SILVA 16S reference database (version 138.2, Chuvochina et al. 2026). We focused on taxonomic classifications at the genus level. We filtered for genera with more than 10 counts across samples and present in at least 4 individuals, resulting in 37 genera represented across samples. Relative abundances of these 37 genera across all samples are listed in Supplementary File 2. We constructed PCA plots of pup samples using the prcomp function in R/stats, and quantified differentially abundant taxa between strains and rearing dam groups using R/DESeq2 (Love et al. 2014) with the design ∼ pop + rearingDam. Genera were considered significantly differentially abundant at a Benjamini-Hochberg adjusted p-value of 0.05.

To quantify variation in alpha and beta diversity metrics across samples, we calculated Shannon’s and Simpson’s indices using the diversity function in R/vegan and calculated Bray-Curtis dissimilarity for all pairwise pup comparisons using the adonis2 function in R/vegan. For Shannon’s and Simpson’s indices, we tested for significant effects of strain and rearing dam using ANOVAs in R/stats. For Bray-Curtis dissimilarity, we conducted pairwise tests of significance using PERMANOVAs in R/pairwiseAdonis.

### Body weight QTL analyses

To identify differential expression linked to phenotypic differences in body weight, we intersected *cis*-DEGs and genes with known phenotypic effects on body size with a published QTL experiment in SARA and MANA mice (Durkin et al. 2026).

First, we tested for enrichment of genes with a known phenotypic effect on body size curated from the Mouse Genome Informatics (MGI) database (Baldarelli et al. 2024) within body weight QTL regions. Specifically, we used the Mammalian Phenotype (MP) code of “abnormal body size” (MP: 0003956), which contains 3,194 genes. We used permutations to test for enrichment of MP genes within genomic regions above the LOD score significance threshold for body weight as determined in Durkin et al. (2026). We linked each MP gene to its closest marker in the QTL map and counted the number of MP genes above the LOD threshold. We then randomly sampled markers of the same size 10,000 times and calculated how many markers were above the LOD threshold in each permutation. We calculated a p-value as the proportion of permuted gene sets that had more markers above the LOD threshold than the MP gene set.

To isolate strong candidate genes, we looked at the intersection between 1) genes above the QTL LOD threshold, 2) MP body size genes, 3) genes with significant *cis*-regulation at 10-days (*cis*-only, *cis* x *trans*, and *cis* + *trans*), 4) genes with significant *cis-*regulation in adult mice from Ballinger et al. (2023), and 5) genes showing evidence of positive selection in wild populations of house mice. For the final criteria, we used published results of a normalized version of the population branch statistic (PBSn1) from Durkin et al. (2024). The PBS test identifies allele frequency shifts that are present in one focal population as compared to two outgroups, and are therefore candidates for loci under selection (Yi et al. 2010; Crawford et al. 2017). We used two published PBSn1 tests, one where a population from Manaus, Brazil is the focal group, and one where a population from the border between New Hampshire and Vermont is the focal group, which is geographically close to Saratoga Springs, the original locality of the SARA line. In both tests, Iran and France were the outgroup populations, which are near the ancestral range of *Mus musculus domesticus* (Phifer-Rixey and Nachman 2015; Ferris et al. 2021; Morgan et al. 2022). We used genes within the top 5% and 10% of PBSn1 scores to intersect with our QTL and expression data.

### Gene ontology enrichment analysis

We used PANTHER (Mi et al. 2013) to identify enriched biological processes among 1) hub genes in the strongly maternal blue module (29 genes) and 2) genes that were *cis*-regulated, above the LOD score significance threshold for body weight, and overlapping a body size MP gene (32 genes). We utilized PANTHER’s hierarchical clustering of GO terms to select the most specific term within a cluster for plotting. Importantly, identifying significant GO terms associated with *cis*-regulated candidate genes is not due to them being a subset of body size MP genes, because 1,000 iterations of random subsets of MP genes of the same size do not show significant enrichments of GO terms to the same degree (Supplementary Figure 8).

### Statistical analyses

Pairwise t-tests throughout are Welch two sample t-tests performed using R/stats.

Additionally, we used linear models to analyze body weight variation of pups and dams during the nursing period while controlling for litter size as a covariate. Estimated marginal means and pairwise comparisons were performed using R/emmeans. Results are in Supplementary Files 6 and 7.

## Resource availability

All sequencing reads related to this study are available on the NCBI Sequence Read Archive under accession IDs PRJNA1480043 (RNA-seq libraries) and PRJNA1482167 (microbiome 16S rRNA ONT libraries). Count matrices and all code used to process and analyze RNA-seq and microbiome data are available on GitHub (https://github.com/s-durkin/weightGainPlasticity_2026). Other data are within the paper and Supplementary Material files.

## Acknowledgements

The authors are grateful to the help of many scientists: Muer Zhou for help in collecting maternal behavioral data; Lydia Smith for guidance during RNA library preparation; David Manahan for assistance in ONT sequencing; and Peter Sudmant for sharing reagents related to 16S rRNA sequencing.

## Author contributions

Conceptualization, S.M.D and M.W.N.; investigation, S.M.D. and C.G.; formal analysis, S.M.D.; software, S.M.D.; writing – original draft, S.M.D.; writing – review and editing, S.M.D., C.G., and M.W.N.; funding acquisition, M.W.N.

