## Supplementary Appendix 1 for "A maternal effect shapes early-life adaptive body size variation in house mice"

**Supplementary Figures**

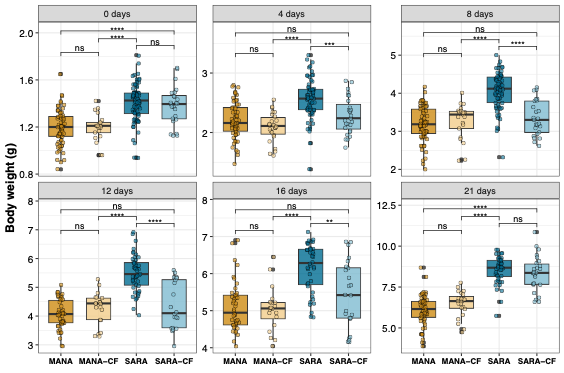

**Supplementary Figure 1: Body weight variation for each timepoint during nursing.** P-values represent Welch two sample t-tests (ns: not significant, * p < 0.05, ** p < 0.01, *** p< 0.001, **** p < 0.0001).

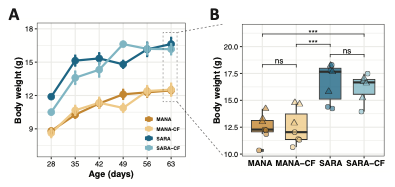

**Supplementary Figure 2: Effect of cross-fostering on body weight into adulthood.** A). Average body weight from four to nine weeks of age. B). Week nine body weight of each pup group. Circles: females, triangles: males. P-values represent Welch two sample t-tests (ns: not significant, * p < 0.05, ** p < 0.01, *** p< 0.001).

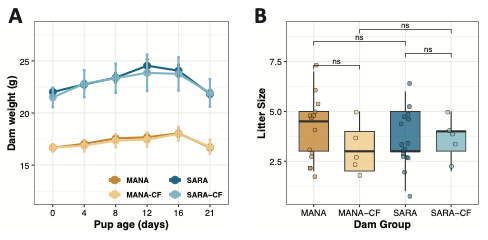

**Supplementary Figure 3: Maternal variation in weight gain and litter size.** MANA-CF and SARA-CF refers to MANA dams rearing SARA litters and SARA dams rearing MANA litters, respectively. A). Dam weight throughout nursing. There are significant differences between SARA and MANA dams at all timepoints, and no differences at any timepoint between dams from the same strain rearing their own litter or a litter of the other strain. Details on statistical tests can be found in Supplementary File 7. B). Litter sizes of each dam group. P-values represent Welch two sample t-tests (ns: not significant).

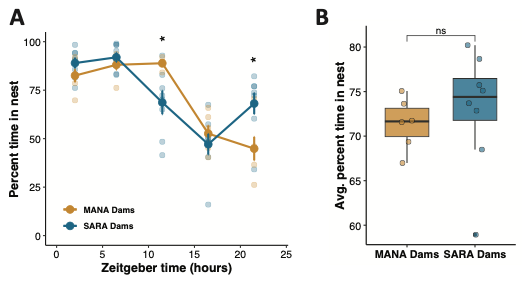

**Supplementary Figure 4: MANA and SARA mothers partition time in the nest differently but spend the same total time in nest throughout the day.** A) Percent of time dams were recorded “in nest” at five timepoints throughout the day. The time in nest averaged across the 6-day recording period is plotted for each separate timepoint. Timepoints without an asterisk are not significantly different. B) Average percent of time in nest across all time points throughout the day. Each dot in A and B represents one individual (MANA dams: n=6, SARA dams: n=8). P-values represent Welch two sample t-tests (ns: not significant, * p < 0.05).

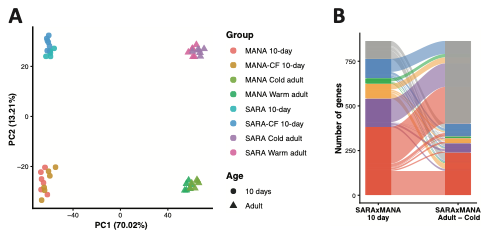

**Supplementary Figure 5. Supplementary material for adult liver gene-regulatory analyses.** A). PCA of all samples from the present study and from a study focusing on adult liver gene expression in MANA and SARA mice (Ballinger et al. 2023). Ballinger et al. (2023) studied gene regulatory patterns in the liver of mice reared at room temperature, and in a cold room (4°C). Only males are included in these analyses. B). Regulatory type alluvial plot of 10-day SARA x MANA males compared to adult SARA x MANA males in the cold condition.

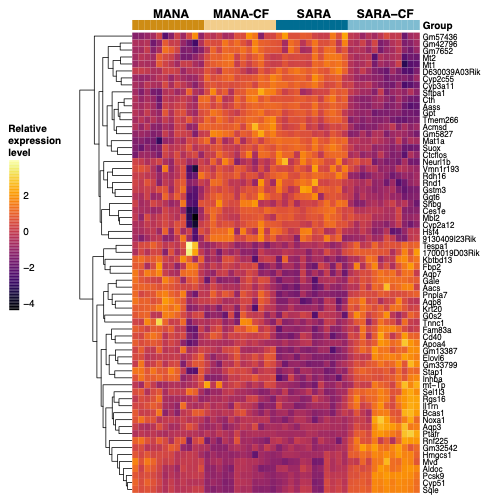

**Supplementary Figure 6: Relative expression level for all genes in maternal module.** WGCNA module most strongly associated with maternal status. Only hub genes are shown in Figure 3.

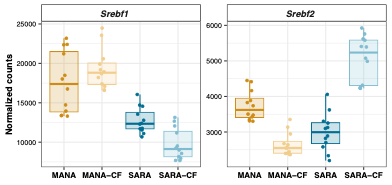

**Supplementary Figure 7: Expression levels of *Srebf1* and *Srebf2*.** Genes encoding sterol regulation element binding proteins are involved in fasting-induced cholesterol pathway activation and have opposing expression patterns in SARA-CF mice.

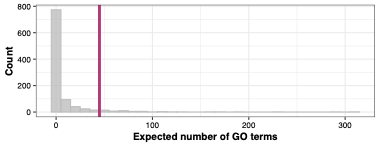

**Supplementary Figure 8: Simulated GO term enrichment among abnormal body size MP code genes.** Grey bars represent the number of GO terms returned from 1,000 iterations of randomly sampling 32 body size MP genes and running a GO term enrichment analysis. The magenta line represents the actual number of GO terms returned (45 terms) when running an enrichment analysis on the 32 *cis*-regulated MP code genes from Figure 5. There are more GO terms in this analysis than that shown in Figure 5C because when plotting, we used Panther’s hierarchical clustering to only display the most specific GO term within each nested category (details in methods). The proportion of the distribution above the line is 0.05, suggesting that the GO terms associated with the 32 *cis*-regulated MP code genes of interest are not purely a function of them being a subset of MP genes.

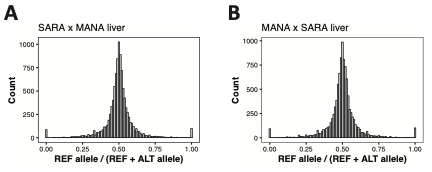

**Supplementary Figure 9: Lack of mapping bias in F1 hybrids.** The ratio of F1 hybrid reads mapping to the MANA allele vs. all mapped reads is centered around 0.5 for SARA x MANA (A) and MANA x SARA (B) F1 hybrids, suggesting there is no mapping bias for reads preferentially mapping to the allele of one parent. REF allele in this context is MANA.

**Supplementary Tables**

**Supplementary Table 1. Significantly differentially expressed genes between male and female MANA non-CF mice.** Significantly differentially expressed genes have an adjusted p-value < 0.05 and a log_2_ fold change > 0.5 or < -0.5.

| Gene ID | Gene symbol | Chr. | Base mean expression | Log_2_ fold change | Adjusted p-value |
| --- | --- | --- | --- | --- | --- |
| ENSMUSG00000025332 | **Kdm5c** | **X** | 3128.85 | -0.48023 | 0.05775 |
| ENSMUSG00000025538 | **Sumf2** | **5** | 243.7947 | 0.61485 | 0.136868 |
| ENSMUSG00000035150 | **Eif2s3x** | **X** | 1769.23 | -0.736 | 0.132031 |
| ENSMUSG00000037369 | **Kdm6a** | **X** | 868.6387 | -0.71194 | 0.118786 |
| ENSMUSG00000056673 | **Kdm5d** | **Y** | 386.752 | 12.00755 | 0.838818 |
| ENSMUSG00000068457 | **Uty** | **Y** | 337.6409 | 11.81176 | 0.838644 |
| ENSMUSG00000069045 | **Ddx3y** | **Y** | 1599.622 | 14.05574 | 0.841514 |
| ENSMUSG00000069049 | **Eif2s3y** | **Y** | 920.6001 | 12.77746 | 0.838088 |
| ENSMUSG00000086370 | **Ftx** | **X** | 173.0131 | -0.7617 | 0.133151 |
| ENSMUSG00000086503 | **Xist** | **X** | 12991.77 | -12.6428 | 0.355255 |
| ENSMUSG00000087174 | **5530601H04Rik** | **X** | 215.6199 | -0.6869 | 0.107321 |
| ENSMUSG00000095041 | **unclassified** | **unc.** | 1656.055 | -0.40007 | 0.058132 |
| ENSMUSG00000097571 | **Jpx** | **X** | 364.6536 | -0.71477 | 0.136738 |
| ENSMUSG00000099876 | **Gm29650** | **Y** | 32.79085 | 8.445242 | 0.862826 |

**Supplementary Table 2. Significantly differentially expressed genes between male and female SARA non-CF mice.** Significantly differentially expressed genes have an adjusted p-value < 0.05 and a log_2_ fold change > 0.5 or < -0.5.

| Gene ID | Gene symbol | Chr. | Base mean expression | Log_2_ fold change | Adjusted p-value |
| --- | --- | --- | --- | --- | --- |
| ENSMUSG00000020038 | **Cry1** | **10** | 838.9848 | 0.472208 | 0.104277 |
| ENSMUSG00000025332 | **Kdm5c** | **X** | 2777.064 | -0.50061 | 0.059386 |
| ENSMUSG00000031226 | **Pbdc1** | **X** | 560.6188 | -0.41404 | 0.066201 |
| ENSMUSG00000034591 | **Slc41a2** | **10** | 293.0181 | 0.436949 | 0.10414 |
| ENSMUSG00000035150 | **Eif2s3x** | **X** | 1892.577 | -0.56258 | 0.073188 |
| ENSMUSG00000037369 | **Kdm6a** | **X** | 852.5389 | -0.6052 | 0.101483 |
| ENSMUSG00000056673 | **Kdm5d** | **Y** | 378.6761 | 11.97964 | 0.835974 |
| ENSMUSG00000068457 | **Uty** | **Y** | 328.2028 | 11.77287 | 0.83998 |
| ENSMUSG00000069045 | **Ddx3y** | **Y** | 1742.582 | 14.18143 | 0.835659 |
| ENSMUSG00000069049 | **Eif2s3y** | **Y** | 872.2767 | 13.18271 | 0.834406 |
| ENSMUSG00000086503 | **Xist** | **X** | 12921.4 | -12.897 | 0.334188 |
| ENSMUSG00000087174 | **5530601H04Rik** | **X** | 177.1203 | -0.48223 | 0.101778 |
| ENSMUSG00000096768 | **Erdr1y** | **Y** | 996.005 | 0.693827 | 0.13404 |
| ENSMUSG00000097571 | **Jpx** | **X** | 306.7039 | -0.66072 | 0.086535 |
| ENSMUSG00000099876 | **Gm29650** | **Y** | 27.05473 | 8.170477 | 0.850146 |

**Supplementary File Captions**

**Supplementary File 1.** Expression levels, fold changes, and differential expression statistics for genetic, MANA maternal, and SARA maternal genes.

**Supplementary File 2.** Relative abundances of genera detected in 16s rRNA gut microbiome samples from MANA and SARA dams and pups.

**Supplementary File 3.** *cis*-DEGs in the liver of 10-day old mice within body weight QTL intervals that overlap mammalian phenotype (MP) body size genes (32 genes total).

**Supplementary File 4.** All significant gene ontology (GO) terms associated with the 32 *cis*-DEG x QTL x MP candidate genes (listed in Supplementary File 3).

**Supplementary File 5.** Genes falling into different regulatory categories for all F1 comparisons. Expression levels, fold changes, and differential expression statistics for parental differential expression, F1 allele differential expression, and *trans*-expression tests are listed.

**Supplementary File 6.** Estimated marginal means and pairwise comparison statistics for pup body weight variation through nursing using litter size as a covariate.

**Supplementary File 7.** Estimated marginal means and pairwise comparison statistics for dam body weight variation through nursing using litter size as a covariate.
